# Programmed death-ligand 1 expression in human cancer three-dimensional cell culture models

**DOI:** 10.1101/2022.10.31.514495

**Authors:** K Hudson, NA Cross, N Jordan-Mahy, R Leyland

**Affiliations:** Biomolecular Sciences Research Centre, Sheffield Hallam University, Sheffield, S1 1WB, UK

**Keywords:** Programmed death-ligand 1, 3D cell culture models, PD-1/PD-L1-targeted therapy, Tumour microenvironment

## Abstract

Programmed death-ligand 1 (PD-L1) expression is a survival mechanism employed by tumours to mediate immune evasion and tumour progression. PD-1/PD-L1-targeted therapies have revolutionised the cancer therapy landscape due to their ability to promote durable anti-tumour immune responses in select patients with advanced cancers. However, some patients are unresponsive, hyper-progressive or develop resistance. Better characterisation of the 3D architecture of solid tumours by utilising 3D cell culture could provide an environment that more closely recapitulates *in vivo* human tumours for investigating tumour-intrinsic PD-L1 signalling and immunotherapy responses. Here we investigated whether PD-L1 expression by human breast, prostate and colorectal cancer cell lines altered in 3D cell culture models compared to their 2D monolayer counterparts. We found that PD-L1 expression changed in 3D-cultured cancer cells when compared to 2D-cultured cells. Additionally, the expression of immunological markers, PD-1, PD-L2, CD44, DR4, DR5, Fas, and HLA-ABC were assessed in 3D cell culture and compared to their expression in 2D. These markers were also altered in 3D compared to 2D-cultured cells, highlighting the importance of utilising 3D models which may better able the investigation of tumour-intrinsic PD-L1 signalling, responses to PD-1/PD-L1-targeted therapy, and combination therapies.

## Introduction

Programmed death-ligand 1 (PD-L1), also known as B7-H1 and CD274, is an immune checkpoint inhibitor expressed by T cells, B cells, natural killer (NK) cells, dendritic cells, macrophages, myeloid-derived suppressor cells, and many other cell types such as epithelial and endothelial cells (Johnson *et al*., 2017; Dong *et al*., 2019). PD-L1 binds to its receptor PD-1 expressed by T cells and less commonly by B cells, activated monocytes, NK cells and dendritic cells to regulate immune responses; ultimately preventing exacerbated activation and autoimmunity (Freeman *et al*., 2000; Dong *et al*., 2002; Jiang *et al*., 2019). During cancer development, anti-tumour immunity is supressed (Hanahan and Weinberg, 2011) and one mechanism in which tumours supress the immune system’s ability to eradicate the tumour is by exploiting the PD-1/PD-L1 signalling axis through overexpression of PD-L1 (Hino *et al*., 2010; Maine *et al*., 2013; Muenst *et al*., 2014; Wang *et al*., 2016). More recently, tumours have also been shown to express PD-1 (Yao *et al*., 2018; Wang *et al*., 2020). Indeed, elevated expression of PD-L1 on tumours has been reported to strongly correlate with advanced disease state and poor prognosis in many different cancers (Wang *et al*., 2016; Hudson *et al*., 2020).

Immunotherapies inhibiting the PD-1/PD-L1 signalling axis have yielded remarkable anti-tumour immune responses in select patients with advanced cancers that present with PD-L1 positive tumours, circulating/intratumoral PD-1 positive CD8 T cells and/or tumours with high mutational burden, and for this reason have become the first line treatment for many cancers (Fehrenbacher *et al*., 2016; Rosenberg *et al*., 2016; Balar *et al*., 2017; Chen *et al*., 2017). However, satisfactory responses to PD-1/PD-L1-targeted therapies are only observed in some patients (∼20-40%) and the reasons for this lack of response and the development of resistance remains poorly understood. The new and emerging roles of both PD-L1 and its receptor PD-1 to send pro-survival signals in tumour cells may be in part responsible for the lack of response to PD-1/PD-L1-targeted therapies, and requires further investigation (Hudson *et al*., 2020). Whilst PD-1/PD-L1-targeted therapies in combination with other immune approaches, small molecule inhibitors, chemotherapy and other modalities are promising strategies to overcome resistance and broaden patient responses (Wang *et al*., 2019), there is still a lack of understanding about how tumorigenic expression of PD-L1 and PD-1 contributes to intrinsic signalling in different cancers. Furthermore, the mechanisms by which anti-PD-1/PD-L1 monoclonal antibodies exert their therapeutic effects on tumours, and whether they can efficiently block tumour-intrinsic signalling of PD-L1 and its receptor PD-1, still requires investigation. Most research investigating the tumour intrinsic roles of PD-L1 and its receptor PD-1 and their response to therapy has been done using monolayer 2D-cultured mouse or human cancer cell lines and/or immunocompromised *in vivo* mouse models. These systems have failed to fully recapitulate *in vivo* human tumours, in terms of their lack of heterogeneity and species-to-species variability (Hudson *et al*., 2020). Hence, there is an urgent need for more relevant *in vitro* oncology models, capable of closely mimicking the heterogeneity of the tumour microenvironment during *in vivo* conditions. This could allow a more predictive *in vitro* evaluation of the tumour-intrinsic role of PD-L1 and its receptor PD-1, and their response to PD-1/PD-L1-targeted therapy, plus enable the evaluation of potential combination strategies and cancer cell-immune cell interactions.

Scaffold-free and scaffold-based 3D cell culture models created with human cancer cells are advantageous alternatives to standard 2D cell culture and *in vivo* mouse models, which are currently the main strategies used as the scientific basis for immuno-oncology studies (Figure 1) (Hudson *et al*., 2020; Boucherit *et al*., 2020). Furthermore, 3D *in vitro* and *ex vivo* (patient-derived) models present with similar biological characteristics to native tumour tissue, having a comparable 3D architecture with a nutrient gradient compromising a proliferative outer layer, a quiescent inner layer and a necrotic core; a hypoxic microenvironment; cell-cell and cell-extracellular matrix signalling; genomic and protein alterations; and a predictive response to treatment (Breslin and O’Driscoll, 2013; Knight and Przybors, 2015; Lazzari *et al*., 2017; Hoarau-Véchot *et al*., 2018; Di Modugno *et al*., 2019; Boucherit *et al*., 2020). These 3D cell culture models also allow the opportunity to alter the cell composition in order to study different aspects of the tumour microenvironment (Boucherit *et al*., 2020).

**Figure 1.**
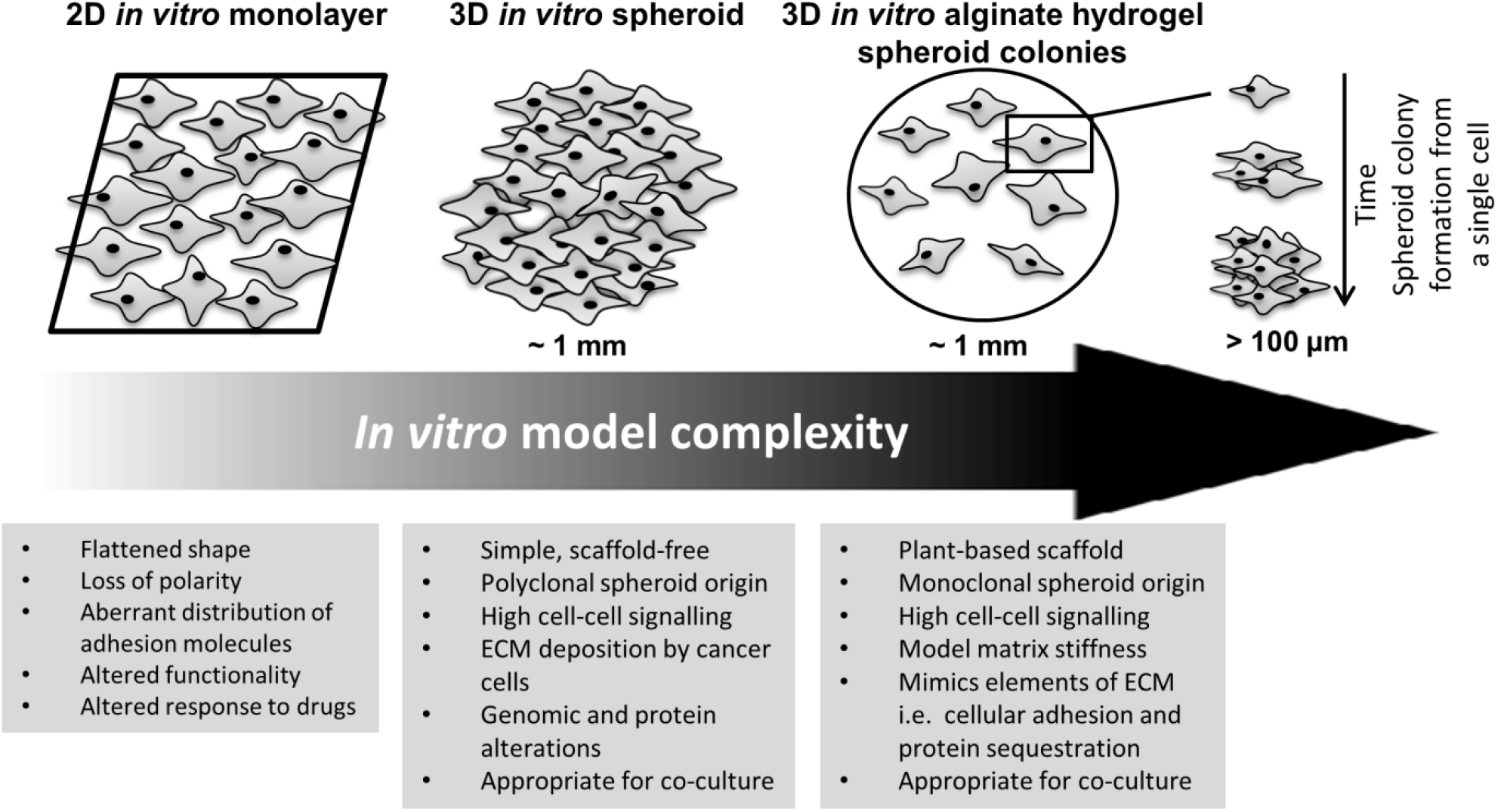
A diagram illustrating the increase in *in vitro* model complexity and the physiological relevance of using these 3D models as opposed to standard 2D monolayer culture for oncology and immuno-oncology studies.

Here we report the expression of PD-L1 and other immunological markers in 3D cell culture models of human breast, prostate and colorectal cancer cell lines compared to their 2D monolayer counterparts. Using standard 2D cell culture, we investigated the mRNA expression and protein production of baseline PD-L1 and the effect of IFNγ and TNFα on cell surface PD-L1 protein expression. Following generation of 3D cell culture models that facilitate the formation of spheroids, we investigated cell viability, mRNA and protein expression of PD-L1 in these 3D models compared to standard 2D cell culture. A subsequent study was made to examine the expression of other immunological markers PD-1, PD-L2, CD44, DR4, DR5 at mRNA and protein levels in 3D cell culture models as well as HLA-ABC cell surface protein expression compared to standard 2D cell culture.

## Materials and methods

### Cell lines and culture conditions

Human breast (MDA-MB-231 and MCF-7), prostate (LNCaP and PC3) and colorectal (SW480 and SW620) cancer cell lines were purchased from American Type Culture Collection (ATCC) (LGC, Teddington, UK) and were cultured in Roswell Park Memorial Institute (RPMI 1640) medium or Dulbecco’s modified Eagle’s medium (DMEM), supplemented with 10% foetal bovine serum (FBS) and 1% penicillin-streptomycin. Cells were maintained in standard culture conditions (5% CO_2,_ 37°C) and grown to 80% confluency before being used experimentally. For each study, the cell concentration of live cells was determined using a haemocytometer and trypan blue stain. All cell culture materials were supplied by Gibco™ (ThermoFisher Scientific, Leicestershire, UK). Cell lines were periodically tested for mycoplasma using the EZ-PCR™ Mycoplasma Detection Kit (Biological Industries, Cromwell, CT, USA) and the MycoAlert™ Mycoplasma Detection Kit (Lonza, Basal, Switzerland) and were confirmed to be mycoplasma free. Cells were below passage 30 for all experiments.

### 2D and 3D cell culture

2D cultures were established in flat-bottom 6-well plates at a seeding density of 500,000 cells/well for each cell line. 3D cultures were established using the hanging drop method (Knight and Przybors, 2015) and alginate hydrogel beads (Arhoma *et al*., 2017).

### 3D hanging drop method

To generate 3D spheroids using the hanging drop method, the lid of a Petri dish was inverted and 10 μL of cell suspension containing 10,000 cells was pipetted drop-by-drop onto the lid. The bottom of the petri dish was filled with PBS to act as a hydration chamber to which the inverted lid was placed on top. The droplets were monitored daily for spheroid formation and were harvested for downstream analysis at day 3, by pipetting 10 mL of complete medium gently onto the lid and collecting all the spheroids, with a single spheroid being generated per droplet.

### 3D alginate hydrogel beads

To form 3D spheroid colonies using alginate hydrogel beads, cell lines were suspended in 1.2% w/v sterile sodium alginate (Merck, Dorset, UK) in 0.15 M sodium chloride at densities ranging from 6 × 10^5^ cells/ml to 1.2 × 10^6^ cells/ml. The alginate-cell mixture was extruded out of a 21G needle into 0.2 M calcium chloride to polymerise the alginate into beads; encapsulating the cells. Following a 10 minute incubation at 37°C alginate hydrogel beads were washed twice with 0.15 M sodium chloride and once with complete medium before culturing for 3, 6 and 10 days. For long term cultures media was changed every 72 hours. Alginate hydrogel beads were monitored daily for spheroid colony formation and harvested at day 3, 6 and 10 for downstream analysis. To release the spheroid colonies into solution, alginate hydrogel beads were immersed in sterile alginate dissolving buffer (55 mM sodium citrate, 30 mM EDTA and 0.15 M sodium chloride) for 10 minutes at 37°C. Spheroid colonies were then washed with PBS in preparation for downstream analysis including real-time quantitative polymerase chain reaction (RT-qPCR) and flow cytometry.

### Monitoring 3D cell culture viability

A single spheroid or a whole alginate hydrogel bead for each cell line was harvested and placed in a 96-well plate and labelled with Hoechst 33342 (10 μg/mL) (ThermoFisher Scientific) and Propidium Iodide (PI) (10 μg/mL) (Merck) for 20 minutes at 37°C. Fluorescent images were obtained using the Olympus IX81 microscope (Olympus Corporation, Southend-on-Sea, UK) and multi-fluorescent images were captured by multiple image alignment at 10x magnification using the Olympus cellSens Imaging Software (Olympus Corporation). After capturing the image, the full spheroid or alginate hydrogel bead was selected, and the surface area of Hoechst 33342 and PI was measured concurrently. The percentage viability for each spheroid and alginate hydrogel bead was calculated by dividing the surface area of Hoechst 33342 stained cells by the total surface area of Hoechst 33342- and PI-stained cells then multiplying by 100.

### RNA extraction and quantification, cDNA synthesis and quantitative polymerase chain reaction

Total RNA was extracted from 2D- and 3D-cultured cells using the ReliaPrep™ RNA Miniprep System (Promega, Madison, WI, USA), according to the manufactures protocol. RNA purity and quantity was assessed by using a NanoDrop 1000 (ThermoFisher Scientific). First-strand complementary DNA (cDNA) was generated from 1 μg of total RNA with the High-Capacity RNA-to-cDNA™ Kit (ThermoFisher Scientific). RT-qPCR was performed using TaqMan® Assays in 10 μL reaction mixtures. TaqMan Assay reaction mixtures contained: 5 μL of TaqMan® Fast Advance (Applied Biosystems, Warrington, UK), 2.5 μL nuclease-free H_2_O, 0.5 μL primer-probe (TaqMan Gene Expression Assays, ThermoFisher Scientific) (Table 1) and 2 μL cDNA or water for a no template control. Hypoxanthine guanine phosphoribosyl transferase gene (HPRT1) and TATA-box binding protein (TBP) were used as endogenous controls. All primer-probes were purchased from Life Technologies Limited (ThermoFisher Scientific). The TaqMan qPCR thermal profile consisted of an initial activation step of 10 minutes at 95°C, followed by 40 cycles of 30 seconds denaturation at 95°C and 1 minute annealing at 60°C. RT-qPCR was performed using the QuantStudio 3 Detection System (QuantStudio Design and Analysis Software, Applied Biosystems, Foster City, CA, USA) and relative expression was calculated using the ΔCT method.

**Table 1.**
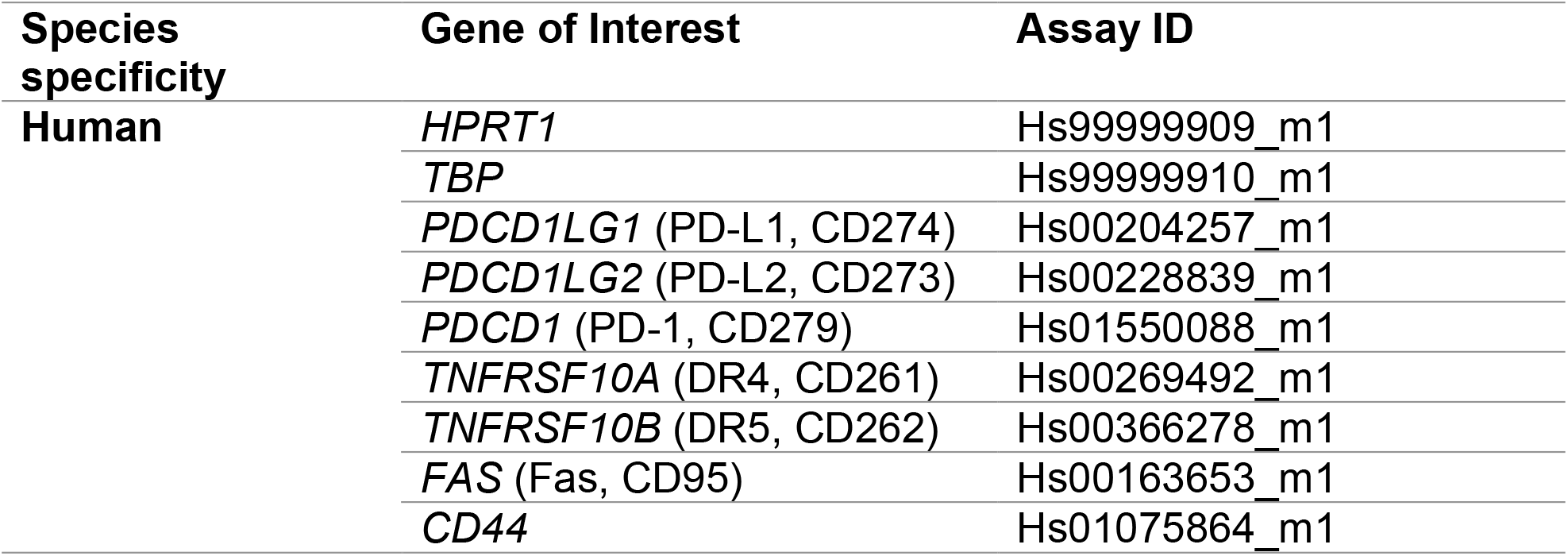
TaqMan primer-probes for RT-qPCR. TaqMan gene expression assay primer-probes (FAM) for RT-qPCR. *HPRT1*, hypoxanthine guanine phosphoribosyl transferase gene; *TBP*, tata box binding protein; *PDCD1LG1* (PD-L1, CD274), programmed death-ligand 1; PDCD1LG2 (PD-L2, CD273), programmed death-ligand 2; *TNFRSF10A* (DR4, CD261), death receptor 4; *TNFRSF10B* (DR5, CD262), death receptor 5; *FAS* (Fas, CD95), Fas; *CD44*, CD44 antigen.

### Flow cytometry

#### Cell surface staining of PD-L1 and other immunological markers

Cells cultured in 2D were dissociated with Trypsin-EDTA 0.25% (ThermoFisher Scientific). Similarly, to achieve single cell suspension for flow cytometry, 3D cultures were dissociated with Trypsin-EDTA 0.25%. 2D and 3D cell suspensions were washed with PBS containing 2% FBS, re-suspended and labelled with Fc block (Human TruStain FcX™; Biolegend, London, UK) for 10 minutes. Cells were subsequently labelled with APC anti-human PD-L1 (clone 29E.2A3; Biolegend), APC anti-human PD-L2 (clone 24F.10C12PE; 2B Scientific Limited, Oxfordshire, UK), anti-human PD-1 (clone: EH12.1; BD Biosciences, San Jose, CA, USA), FITC anti-human HLA-ABC (clone G46-2.6; BD Biosciences), APC anti-human DR4 (clone DJR1; Biolegend), PE anti-human DR5 (clone DJR2-4; Biolegend), PE anti-human Fas (clone DX2; Biolegend) and APC anti-human CD44 (clone IM7; eBiosciences, ThermoFisher Scientific) antibodies or their matched isotype controls for 30 minutes. Data was acquired using either the FACSCalibur (BD Biosciences) with CellQuest™ Pro Software v.5.2.1 (BD Biosciences) or CytoFLEX (Beckman Coulter, IN, USA) with CytExpert Software (Beckman Coulter). Data was analysed using FlowJo 10 Software (FlowJo, LLC, Ashland, OR, USA). Appropriate gating strategies were carried out for each independent experiment including gating on 2D and 3D isotype controls, single cells and live cells. For MDA-MB-231 cells, isotype controls for 2D-cultured cells and 3D alginate spheroid colonies are both displayed on representative flow cytometry plots for DR5 and Fas due to differences in background signal in unstained cells between culture methods. A PE-conjugated isotype control that was a different clone (clone P3.6.2.8.1; eBiosciences) to the original isotype clone used in conjugation with DR5 and Fas antibodies was investigated in 2D-cultured and 3D alginate-cultured cells to exclude non-specific binding as a factor for increased background in 3D isotype controls compared to 2D. The percentage expression and median fluorescent intensity (MFI) of each sample run was determined by dividing the corresponding value by the isotype control.

#### Cytokine modulation of PD-L1 expression

Cells cultured in 2D were seeded at 5 × 10^5^ cells/well in 6-well plates and incubated with recombinant human interferon gamma (IFNγ) (0.5 ng/mL) (Biolegend) and/or recombinant tumour necrosis factor alpha (TNFα) (5 ng/mL) (Bio-Techné, Minneapolis, MN, USA) for 48 hours before assessing the cell surface expression of PD-L1 using flow cytometry.

#### Statistical analysis

Statistical analysis was performed using Prism version 7.03 (GraphPad Software, Inc., La Jolla, CA, USA) and StatsDirect 3 (StatsDirect Ltd, Merseyside, UK). A Shapiro-Wilk normality test was used to determine whether data was normally distributed. For non-parametric data with 2 groups a Mann-Whitney U test was performed, otherwise, Kruskal-Wallis was used followed by the Conover-Iman multiple comparison test. Data shown here represents median ± interquartile range (IQR) and values less than 0.05 were considered significant (*P<0.05, **P<0.01, ***P<0.001 ****P<0.0001). Each independent experiment has 3 technical repeats (n=3).

## Results

### Human breast, prostate and colorectal cancer cell lines express differential levels of PD-L1 at mRNA and protein level in 2D cell culture

Six human cancer cell lines including two breast (MDA-MB-231 and MCF-7), two prostate (LNCaP and PC3) and two colorectal (SW480 and SW620) were assessed to determine the basal level of PD-L1 expression at mRNA and protein level under standard *in vitro* culture conditions using RT-qPCR and flow cytometry, respectively. PD-L1 expression at mRNA (Figure 2A) and protein (Figure 2B and Figure 2C) levels for each human cancer cell line were analysed to show the differential expression across the cancer types.

**Figure 2.**
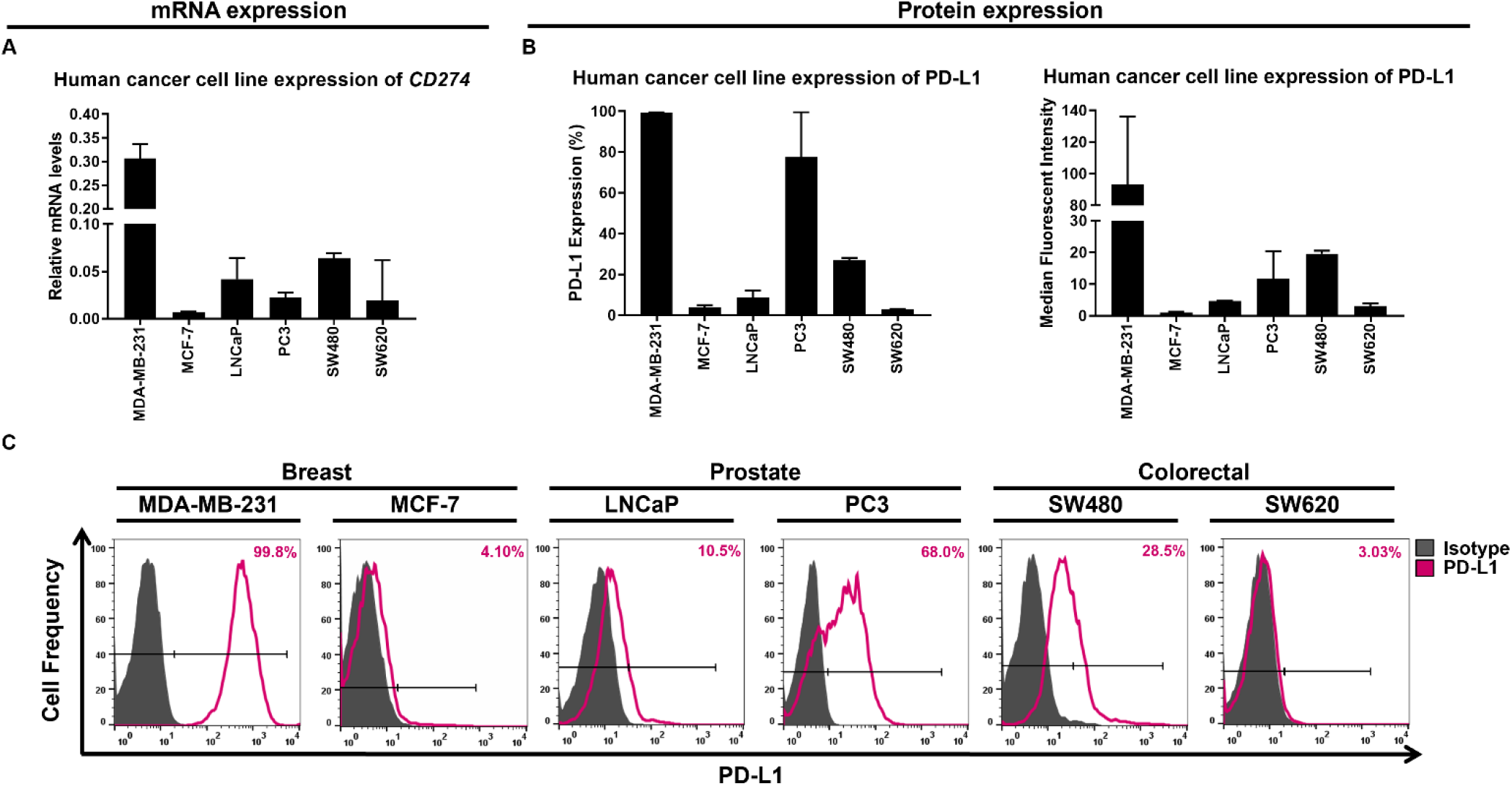
Human breast, prostate and colorectal cancer cell lines express differential levels of PD-L1 at mRNA and protein level in 2D cell culture. PD-L1 (*CD274*) gene expression was measured by RT-qPCR in MDA-MB-231, MCF-7, LNCaP, PC3, SW480 and SW620 cancer cell lines cultured in 2D (**A**). Flow cytometric analysis was used to measure the cell surface expression of PD-L1 in MDA-MB-231, MCF-7, LNCaP, PC3, SW480 and SW620 cancer cell lines cultured in 2D (**B**). The percentage of PD-L1 expression is shown alongside the MFI for each cell line. Representative flow cytometry histograms are displayed for each cancer cell line showing the isotype control (grey) relative to the PD-L1 positive population (pink) (**C**). Date is presented as median ± range, n=3 independent experiments each with 3 technical repeats.

RT-qPCR and flow cytometric data showed that PD-L1 expression was differentially expressed across all human cancer cell lines at mRNA and protein levels. MDA-MB-231 triple negative breast cancer (TNBC) cells expressed the highest level of PD-L1 at both mRNA and protein levels; when compared to cells grown in standard 2D culture conditions. Virtually all MDA-MB-231 cells expressed PD-L1 (99.5% ± 0.159), whereas in MCF-7 luminal-derived, hormone-expressing breast cancer cells expressed one of the lowest levels of PD-L1 at mRNA and protein level; with only a small percentage of cells expressing this low level of PD-L1 (4% ± 1.818). LNCaP lymph node-derived metastatic prostate cancer cells expressed a low to moderate level of PD-L1 at mRNA and protein levels, and like MCF-7 cells, only a small proportion (8.6% ± 8.24) of LNCaP cells expressed a low level of PD-L1 on their cell surface. In comparison, a high proportion of PC3 bone-derived metastatic prostate cancer cells expressed PD-L1 on their cell surface (77.5% ± 31.73), however, the level of PD-L1 expression by PC3 cells was relatively low, which was shown to be consistently low at the mRNA level. For SW480 primary-derived colorectal cancer cells over one quarter of cells expressed PD-L1 on their cell surface (27% ± 1.655) at a moderate level. Whilst the SW480 cells expressed the second highest PD-L1 expression at the mRNA level, although this was still relatively low in comparison to MDA-MB-231 cells. Interestingly, SW620 lymph node-derived metastatic colorectal cancer cells originating from the same tumour of which SW480 cells derived, expressed the lowest frequency of PD-L1 on their cell surface out of all the cell lines with only 3% (± 1.544) of cells expressing PD-L1 at a low level. Metastatic SW620 cells also expressed half the level of PD-L1 mRNA compared to primary SW480 cells.

### IFNγ and TNFα synergistically upregulate cell surface PD-L1 in some human cancer cell lines grown in 2D cell culture

Next, we determined whether PD-L1 protein expression could be modulated on the cell surface of human cancer cell lines cultured under standard 2D culture conditions. Human cancer cells were treated with IFNγ and TNFα either alone or in combination. Our data showed that PD-L1 expression in 4 out of the 6 cancer cell lines could be upregulated by individual cytokines and synergistically upregulated when treated with both cytokines (Figure 3). With only MDA-MB-231 breast (Figure 3A) and LNCaP prostate cancer cells (Figure 3C) showing no significant difference in PD-L1 expression when treated with individual or both cytokines, although these cells did demonstrate a trend increase in PD-L1 expression with cytokine treatment. MDA-MB-231 cells showed a trend of increased PD-L1 expression with individual cytokine treatment, which was further enhanced with combined cytokine treatment, whereas, LNCaP cells demonstrated a trend of increased PD-L1 expression only in response to treatment with TNFα.

**Figure 3.**
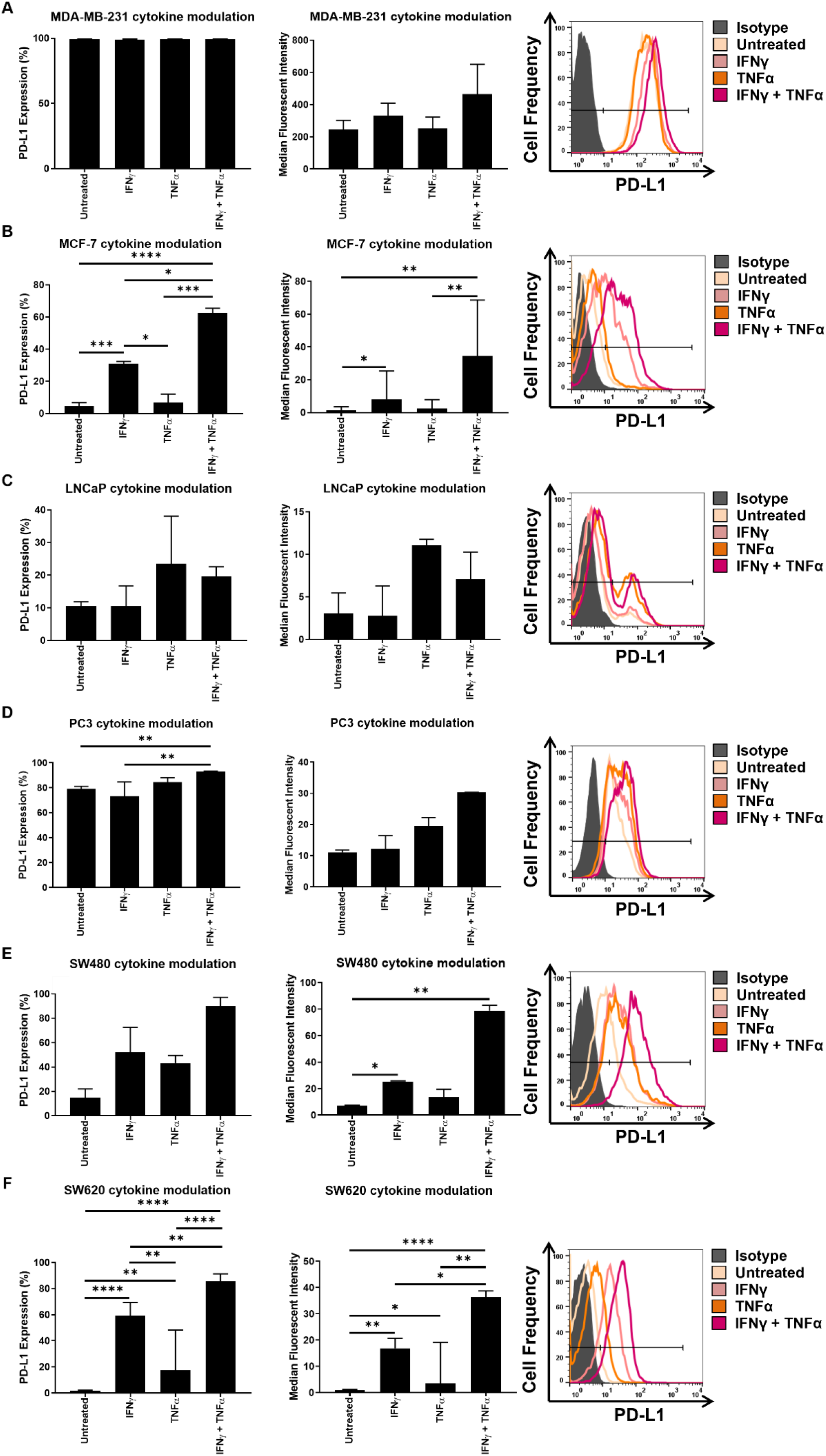
IFNγ and TNFα act synergistically to upregulate PD-L1 expression in human breast MCF-7 cells, prostate PC3 cells and colorectal SW480 and SW620 cells. Flow cytometric analysis was used to determine the effects of IFNγ and/or TNFα on cell surface PD-L1 expressed by six human cancer cell lines. Cancer cells were treated with the lowest significant dose of IFNγ and TNFα alone and combined. The percentage of cells expressing PD-L1 is shown alongside the MFI for each cell line. Representative flow cytometry histograms are displayed showing the isotype control (grey), untreated PD-L1 positive cells (beige), IFNγ treated cells (light pink), TNFα treated cells (orange) and both IFNγ and TNFα treated cells (dark pink). Data is presented as median ± range, n=3 independent experiments each with 3 technical repeats, Kruskal-Wallis followed by Conover-Iman multiples comparison test, *P<0.05, **P<0.01, ***P<0.001, ****P<0.0001.

Furthermore, MCF-7 breast cancer cells displayed a significant increase in the frequency of cells expressing PD-L1 with individual and combined cytokine treatment, as well as a significant increase in the MFI of PD-L1 expression following treatment with IFNγ alone, which was then further enhanced when combined with TNFα compared to the untreated cells (Figure 3B). PC3 prostate cancer cells exhibited a significant increase in the proportion of cells expressing PD-L1, but exhibited no change in the MFI following treatment with both cytokines compared to untreated cells and cells treated with IFNγ and TNFα alone (Figure 3D). SW480 colorectal cancer cells displayed no significant change in frequency of cells expressing PD-L1, but IFNγ alone and in combination with TNFα significantly upregulated the MFI of PD-L1 expression compared to the untreated cells (Figure 3E). SW620 colorectal cancer cells displayed a significant increase in the proportion of cells expressing PD-L1 and MFI when treated with cytokines alone and together compared to the untreated cells (Figure 3F). The frequency and level of PD-L1 expression by SW620 cells significantly increased following treatment with both cytokines compared to individual cytokines alone.

### Hanging drop cancer spheroids display altered PD-L1 expression when compared to cancer cells in 2D cell culture

To mimic more closely the physiological characteristics of solid tumours we used the hanging drop method (Figure 4A) to generate human breast, prostate and colorectal cancer 3D spheroids and assessed these for changes in PD-L1 expression at mRNA and protein level compared to 2D-cultured cells. Here, single spheroids were generated by the hanging drop method and harvested at day 3 to be stained with Hoechst 33342/PI to assess cell viability using fluorescence microscopy. All six human cancer cell lines formed viable and intact spheroids at day 3 (Figure 4B and Figure 4C). The cell viability of 3D spheroids was further validated by Annexin V/PI staining after spheroids were subjected to disaggregation for downstream analysis using flow cytometry (Figure 4D).

**Figure 4.**
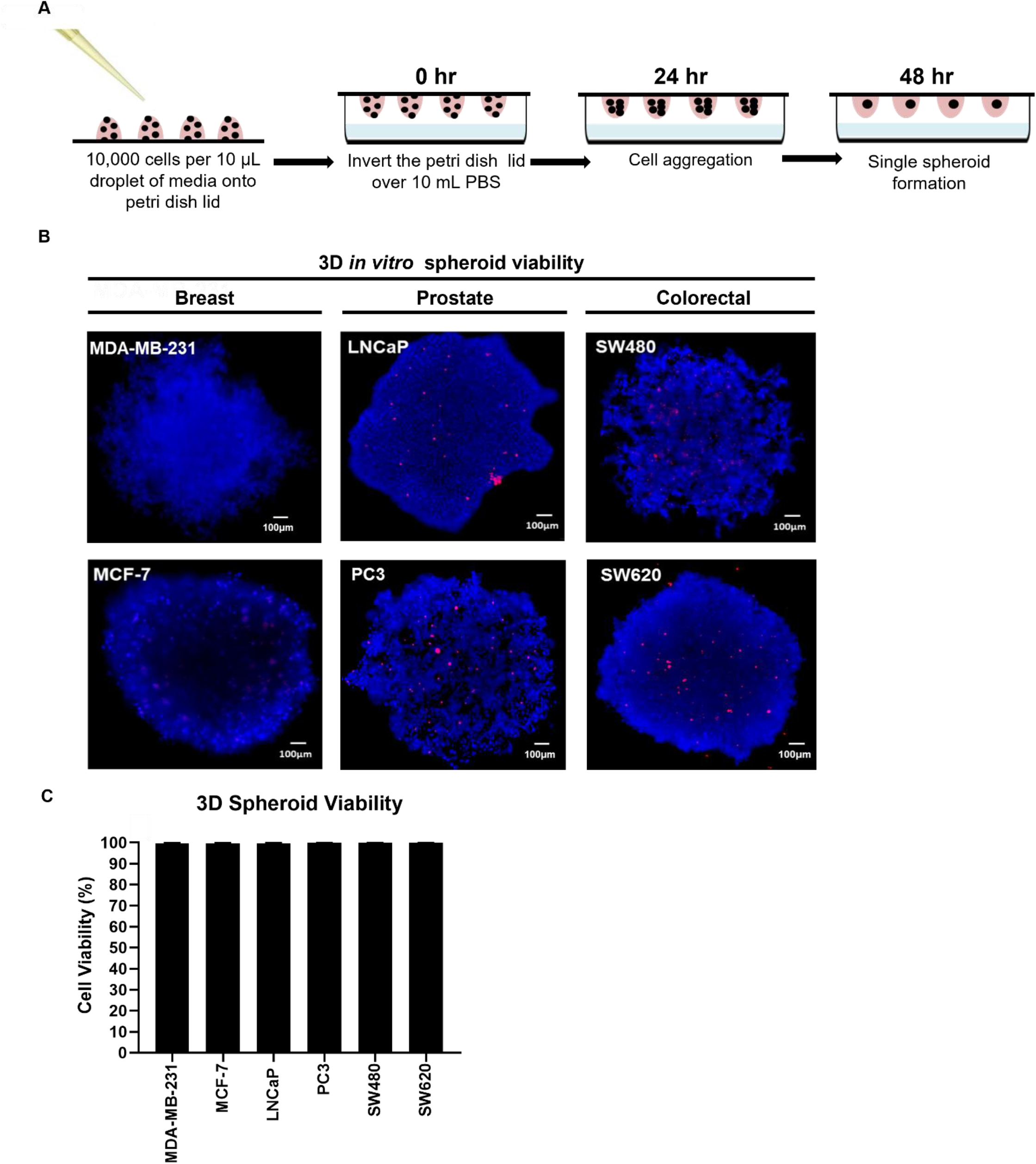
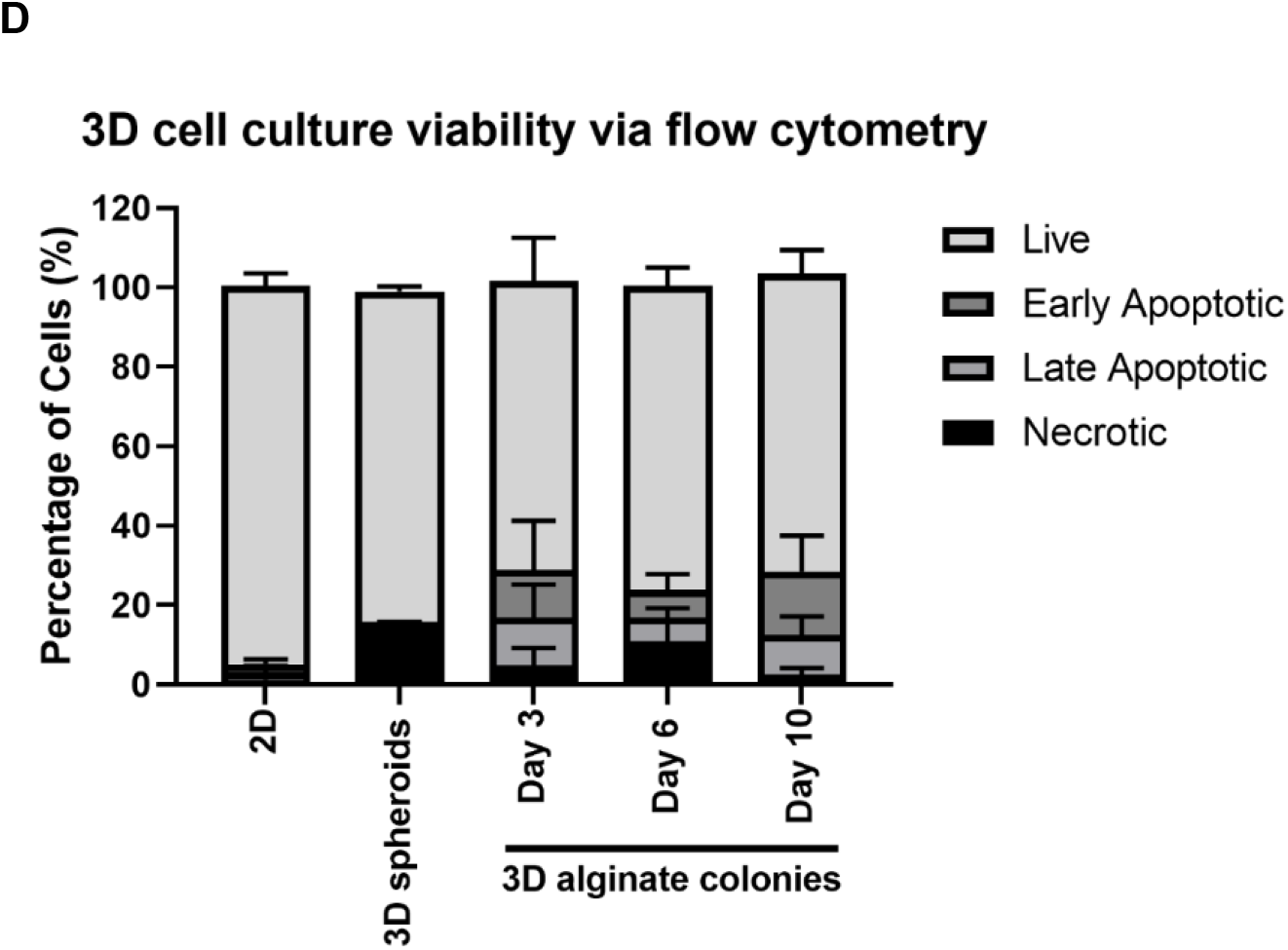
Human breast, prostate and colorectal cancer cell lines form viable and intact 3D spheroids using the hanging drop method. A schematic diagram illustrating the simple steps of the hanging drop method that leads to the formation of a single spheroid per droplet (**A**). Single 3D spheroids generated by the hanging drop method were harvested at day 3 for viability assessment by performing fluorescence microscopy following Hoechst 33342/PI staining. Multiple image alignments of 3D spheroids were captured for each cell line including breast (MDA-MB-231 and MCF-7), prostate (LNCaP and PC3) and colorectal (SW480 and SW620) cancer cells (**B**). Scale bar represents 100 μm. 3D spheroid viability was measured and calculated has a percentage for each cell line (**C**). Data is presented as median ± range, n=3 independent repeats. **(D)** For flow cytometric analysis, 3D cultures were disaggregated to single cell suspensions and the effects of this disaggregation on cell viability was measured by Annexin V/PI staining via flow cytometry. Representative viability data for MDA-MB-231 breast cancer cells Data is presented as median ± range, n=3 independent experiments each with 3 technical repeats.

The expression of PD-L1 at mRNA and protein levels was determined by RT-qPCR and flow cytometry, respectively, for all six human cancer cell lines grown in standard 2D cell culture and 3D hanging drop spheroids. Our data revealed that PD-L1 mRNA and/or protein expression altered in all human cancer cell lines grown in 3D spheroids compared to their 2D counterparts (Figure 5). Appropriate gating strategies were performed including single cell and live cell gating for each experiment (Figure 5A). MDA-MB-231 breast cancer cells exhibited a significant decrease in the level of PD-L1 mRNA (Figure 5B) and protein (Figure 5C) expression in hanging drop 3D cell culture, compared to 2D-cultured cells. MCF-7 breast cancer cells displayed a significant increase in the frequency of cells expressing PD-L1 in 3D cell culture, but the level of PD-L1 mRNA (Figure 5D) and protein (Figure 5E) expression was similar to 2D-cultured cells. LNCaP prostate cancer cells showed no difference in PD-L1 expression at mRNA level (Figure 5F), but did display a significant increase in the proportion of cells expressing PD-L1, and the level of PD-L1 protein on their cell surface in 3D cell culture compared to 2D (Figure 5G). PC3 prostate cancer cells demonstrated a significant decrease in the level of PD-L1 expression at mRNA (Figure 5H) level and in the percentage of cells expressing cell surface PD-L1 (Figure 5I). SW480 colorectal cancer cells exhibited a significant decrease in the mRNA level of PD-L1 (Figure 5J). Interestingly, the frequency of SW480 cells expressing PD-L1 and the level of PD-L1 protein on the cell surface significantly increased in 3D cell culture (Figure 5K). Similar to MCF-7 cells, SW620 colorectal cancer cells exhibit a significant increase in the proportional of cells expressing cell surface PD-L1 in 3D, but the level of PD-L1 expression was similar to 2D at mRNA (Figure 5L) and protein (Figure 5M) levels.

**Figure 5.**
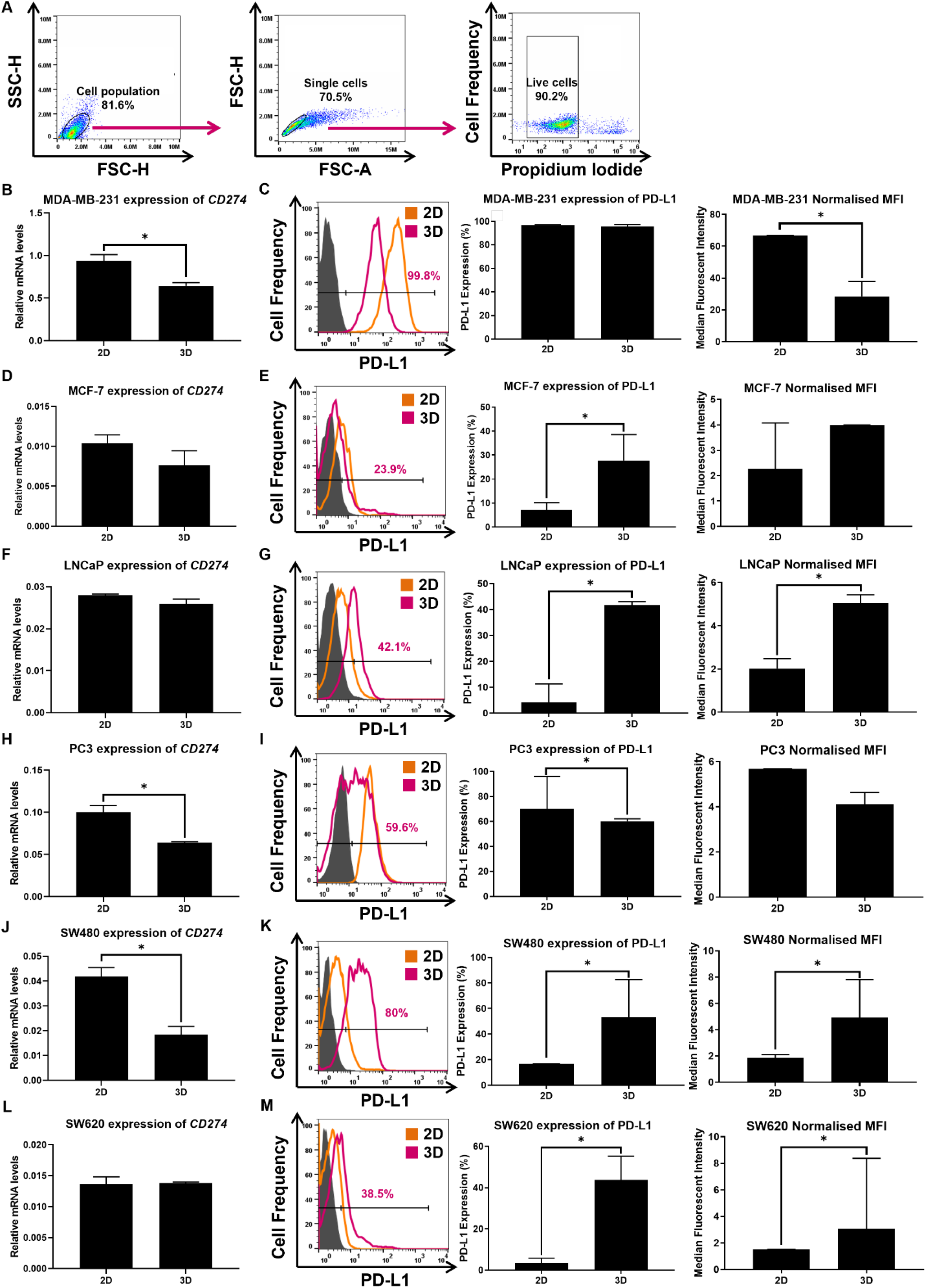
3D cancer spheroids generated using the hanging drop method display altered PD-L1 mRNA and protein expression when compared to cancer cells grown in 2D monolayer cell culture. PD-L1 mRNA and protein expression was measured by qPCR and flow cytometry, respectively, in 2D-cultured cells and 3D spheroids formed using the hanging drop method at day 3. For flow cytometry, appropriate gating strategies including single cell and live cell gating were performed for each experiment **(A)**. PD-L1 mRNA expression in 2D- and 3D-cultured cells was measured for each human cancer cell line including MDA-MB-231 (**B**), MCF-7 (**D**), LNCaP (**F**), PC3 (**H**), SW480 (**J**) and SW620 (**L**) at day 3. Cell surface expression of PD-L1 in 2D- and 3D-cultured cells was measured for each human cancer cell lines including MDA-MB-231 (**C**), MCF-7 (**E**), LNCaP (**G**), PC3 (**I**), SW480 (**K**) and SW620 (**M**) at day 3. Representative flow cytometry histograms are displayed for each cell line showing the isotype control (grey) relative to the PD-L1 positive populations for cells grown in 2D (orange) and 3D (pink) cell culture. The percentage of PD-L1 positive cells is shown alongside the normalised MFI for each cell line. Data is presented as median ± IQR, n=3 independent experiments each with 3 technical repeats, Mann-Whitney U test, *P<0.05.

This data shows that some human cancer cell lines that display a high frequency of cells expressing PD-L1 (MDA-MB-231 and PC3 cells) in standard 2D cell culture, can exhibit a significant decrease in the levels of PD-L1 at mRNA and/or protein levels in 3D cell culture. In comparison, some human cancer cell lines that have a low frequency of cells that express PD-L1 in 2D cell culture display either a significant increase in the frequency of cells expressing PD-L1 in 3D cell culture (MCF-7 and SW620 cells) or exhibit a significant increase in both the frequency and the MFI of PD-L1 expression at protein level in 3D (LNCaP and SW480 cells).

### Cancer 3D spheroid colonies generated in alginate hydrogel beads exhibit altered PD-L1 expression compared to cancer cells in 2D cell culture

To evaluate whether the alterations observed in PD-L1 expression were consistent across different *in vitro* 3D cell culture models, we investigated the expression of PD-L1 at mRNA and protein level in a second, more complex alginate hydrogel bead 3D cell culture model that incorporates a biologically inert scaffold and relies on clonal growth rather than aggregation of cells (Figure 6A). Here, all cancer cell lines grew successfully in the alginate hydrogel beads and formed 3D spheroid colonies reaching ∼100 μm in diameter at day 10 (Figure 6B). Hoechst 33342/PI staining did not reveal areas of pronounced cell death caused by the fabrication procedure. The cells within the alginate remained viable over the 10 day period investigated, displaying an average viability greater than 90% at each time point (Figure 6C). The viability of cells forming 3D spheroid colonies within the alginate were further assessed using Annexin V/PI staining via flow cytometry after dissociation from the alginate and disaggregation necessary for downstream analysis (Figure 4D).

**Figure 6.**
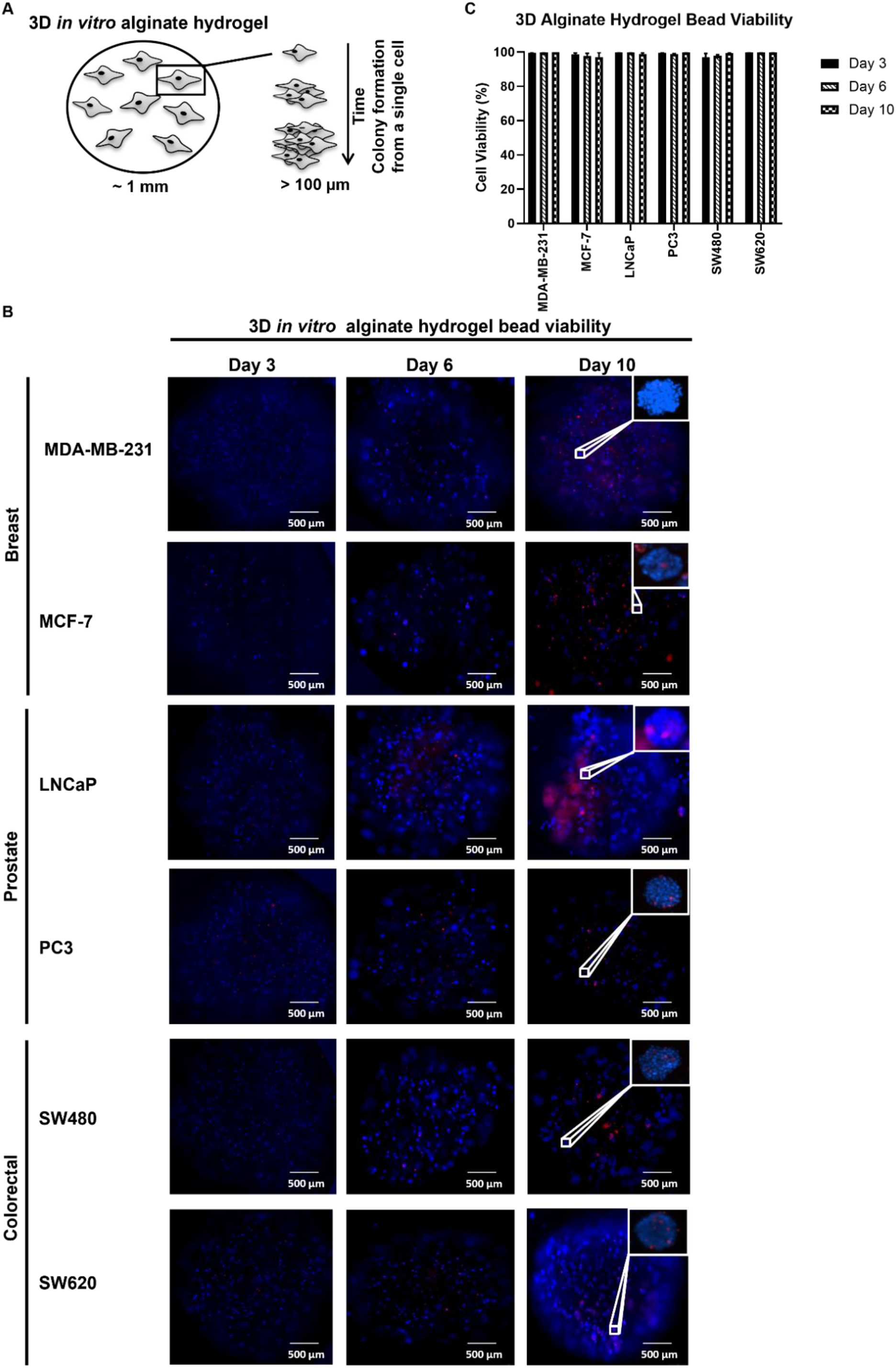
Human breast, prostate and colorectal cancer cell lines grow successfully in alginate hydrogel beads to form monoclonal 3D spheroid colonies of ∼100 μm at day 10. Human cancer cells were encapsulated in alginate hydrogel and grown in culture for up to 10 days by which time individual cancer cells were able to proliferate overtime to form monoclonal spheroid colonies of ∼100 μm (**A**). Cell viability of cancer cells growing in the alginate hydrogel was assessed at day 3, 6 and 10 by fluorescence microscopy via Hoechst/PI staining. For each time point, fluorescent images of whole intact beads are shown for each cell line (**B**). Scale bar represents 500 μm. The images taken at a higher magnification of individual 3D spheroid colonies shown in the white boxes represent spheroid colonies within the alginate hydrogel at day 10 for each cell line that have a diameter of ∼100 μm. At each time point the viability of a whole alginate bead was measured and calculated has a percentage for each cell line (**C**). Data is presented as median ± IQR, n=3 independent repeats.

For 3D spheroid colonies formed in alginate, PD-L1 expression at mRNA and protein level was assessed at day 3, 6 and 10 using RT-qPCR and flow cytometry, respectively. For flow cytometry experiments appropriate gating strategies were performed including single cell and live cell gating for each experiment (Figure 7A). Expression data revealed that the 3D environment created by both 3D cell culture models similarly modulated changes to PD-L1 expression at mRNA and/or protein level in all human cancer cell lines investigated (Figure 5 and Figure 7). It is important to note that these changes in PD-L1 expression in both 3D cell culture models were shown not to be due to disaggregation of 3D cultures to single cell suspensions (Data not shown). Alginate dissolving buffer treatment prior to downstream analysis of PD-L1 expression did show to slightly reduce PD-L1 MFI in MCF-7 and LNCaP cells. Reassuringly, we observe increases in PD-L1 MFI in 3D-cultured MCF-7 and LNCaP cells and therefore have to be mindful that this increase in PD-L1 expression could be higher in these cells in 3D cell culture if it was not necessary to dissolve the alginate in this buffer prior to downstream analysis.

**Figure 7.**
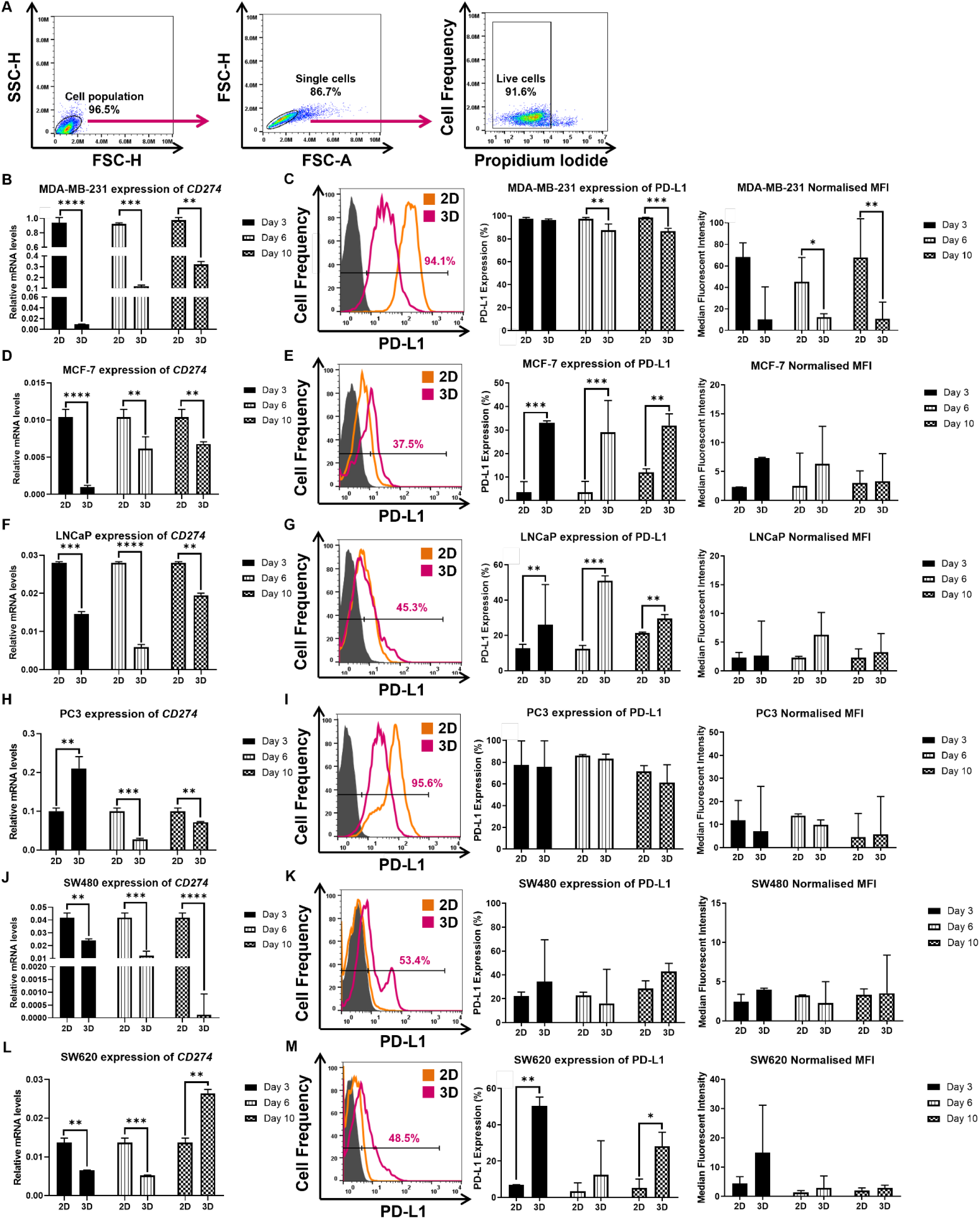
Monoclonal 3D spheroid colonies formed in alginate hydrogel beads exhibit altered PD-L1 mRNA and protein expression when compared to cancer cells grown in 2D monolayer cell culture. 2D-cultured cells and alginate-derived cancer cell 3D spheroid colonies were harvested at day 3, 6 and 10 to measure PD-L1 mRNA and protein expression by RT-qPCR and flow cytometry, respectively. For flow cytometry, appropriate gating strategies including single cell and live cell gating were performed for each experiment **(A)**. PD-L1 mRNA expression in 2D- and 3D-cultured cells was measured for each human cancer cell line including MDA-MB-231 (**B**), MCF-7 (**D**), LNCaP (**F**), PC3 (**H**), SW480 (**J**) and SW620 (**L**) at day 3, 6 and 10. Cell surface expression of PD-L1 in 2D- and 3D-cultured cells was measured for each human cancer cell line including MDA-MB-231 (**C**), MCF-7 (**E**), LNCaP (**G**), PC3 (**I**), SW480 (**K**) and SW620 (**M**) at day 3, 6 and 10. Representative flow cytometry histograms are displayed for each cell line showing the isotype control (grey) relative to the PD-L1 positive populations for cells grown in 2D (orange) and 3D (pink) cell culture. The percentage of cells expressing PD-L1 is shown alongside the normalised MFI for each cell line. Data is presented as median ± IQR, n=3 independent experiments each with 3 technical repeats, Kruskal-Wallis followed by Conover-Iman multiples comparison test, *P<0.05, **P<0.01, ***P<0.001, ****P<0.0001.

MDA-MB-231 breast cancer 3D spheroid colonies grown in alginate displayed a significant decrease in the frequency of cells expressing PD-L1 and the level of PD-L1 expressed at mRNA and protein levels over time, when compared to cells in standard 2D cell culture (Figure 7B and Figure 7C). The significant decrease in the level of PD-L1 expression at mRNA and protein levels observed in the alginate model was consistent with MDA-MB-231 cancer cells grown in 3D spheroids generated by the hanging drop method. MCF-7 breast cancer 3D spheroid colonies showed a significant decrease in the mRNA expression of PD-L1 at each time point (Figure 7D). However, they displayed a significant increase in the frequency of cells expressing cell surface PD-L1 at all time points in 3D in comparison to 2D-cultured cells (Figure 7E). The frequency and level of PD-L1 protein expression by MCF-7 cancer cells grown in alginate was comparable to that observed in cells cultured using the hanging drop method; both 3D culture models showed significantly increased frequency of PD-L1 expression compared to 2D-cultured cells, but the MFI was similar to that seen in 2D-cultured cells. Similarly, LNCaP prostate cancer 3D spheroid colonies grown in alginate showed a significant decrease in PD-L1 mRNA (Figure 7F) and exhibited a significant increase in the frequency of cells expressing PD-L1 at each time point in 3D compared to standard 2D cell culture (Figure 7G). This significant change observed in the percentage of LNCaP cells expressing PD-L1 protein in the alginate model was reciprocated in the hanging drop model, but at a similar MFI to 2D-cultured cells. PC3 prostate cancer cells that displayed a significant decrease in PD-L1 mRNA and protein expression in hanging drop 3D spheroids compared to 2D-cultured cells, also demonstrate a significant decrease in PD-L1 mRNA expression when grown in alginate (Figure 7H). However, in PC3 cells grown in alginate the frequency of PD-L1 expression and the MFI remained unchanged compared to 2D-cultured cells (Figure 7I). Likewise, SW480 colorectal cancer cells grown in alginate displayed a significant decrease in PD-L1 mRNA expression compared to 2D-cultured cells (Figure 7J) as they did in hanging drop 3D spheroids. The frequency of PD-L1 protein and MFI was similar to 2D-cultured cells when SW480 cells were grown in alginate (Figure 7K) which differs from the significant increase in PD-L1 protein observed in SW480 cells cultured using the hanging drop model. Furthermore, SW620 colorectal cancer cells grown in 3D spheroid colonies formed in alginate exhibited a significant decrease in PD-L1 mRNA expression at day 3 and 6 which was followed by a significant increase at day 10 compared to mRNA levels in 2D-cultured cells (Figure 7L). SW620 3D spheroid colonies also showed a significant increase in the frequency of cells expressing PD-L1 compared to 2D-cultured cells at day 10 (Figure 7M). In summary, the level of PD-L1 protein expressed by SW620 cells grown in alginate was similar to that of 2D-cultured cells and 3D spheroids generated by the hanging drop method; only the mRNA levels differed between both 3D models.

This data reveals that human breast, prostate and colorectal cancer cell lines investigated here can be implemented into two different 3D cell culture models and are able to grow viable aggregation-based 3D spheroids and clonal-based 3D spheroid colonies. Moreover, these data show that the expression of PD-L1 by 2D- and 3D-cultured cancer cells can be measured specifically and that consistent change in PD-L1 expression can be detected across two 3D cell culture models in all human cancer cell lines studied.

### Immunological markers expressed by human cancer cells differ in their expression levels in 3D-cultures compared to their 2D monolayer counterparts

Since PD-L1 expression by human cancer cells differed in a 3D cell culture environment, we investigated whether these changes in 3D could be reciprocated for other immunological markers at mRNA and/or protein level. Representative flow cytometry histograms for protein expression of immunological makers by cancer cells grown in 3D hanging drop spheroids (Figure 8) and 3D alginate spheroid colonies (Figure 9) are displayed for each cancer cell line investigated. PD-1, the receptor for PD-L1 and PD-L2, recently been reported to be expressed by tumour cells was investigated here and was found to only be expressed at very low levels in colorectal cancer cell lines (SW480 and SW620) at mRNA level in 2D- and 3D-cultures (Figure 10A). PD-1 expression at protein level was only detected in 1.2% of SW480 cancer cells in 2D culture which significantly increased to 13% in 3D spheroid colonies formed in alginate hydrogel beads (Figure 10B). However, PD-1 protein expression was no longer detectable in 3D spheroids formed by hanging drop. PD-L2 expression was also assessed and was detected in MDA-MB-231, MCF-7, PC3 and SW620 cells at mRNA (Figure 10C) and protein (Figure 10D) level which was found to be expressed at a lower level or similar level in 3D cultures compared to 2D-cultured cells. Interestingly, in MDA-MB-231 and PC3 cancer cells where PD-L1 mRNA expression was found to be significantly decreased in 3D cultures compared to 2D, the expression of PD-L2 was also significantly decreased at mRNA level in 3D cultures compared to 2D-cultured cells. Additionally, when the expression of CD44, a cell surface adhesion receptor was measured, similar trends in its expression were observed amongst cell lines also correlating with PD-L1 expression in 2D and 3D cultures at either mRNA (Figure 10E) and/or protein (Figure 10F) levels. MDA-MB-231 and PC3 cells express lower levels of CD44 mRNA in 3D cell culture compared to 2D-cultured cells; these cells also displayed significantly reduced PD-L1 mRNA in 3D compared to 2D. Conversely, MCF-7, SW480 and SW620 cells that demonstrated a higher proportion of cells expressing PD-L1 in 3D compared to 2D-cultured cells displayed increased CD44 mRNA in 3D culture compared to 2D.

**Figure 8.**
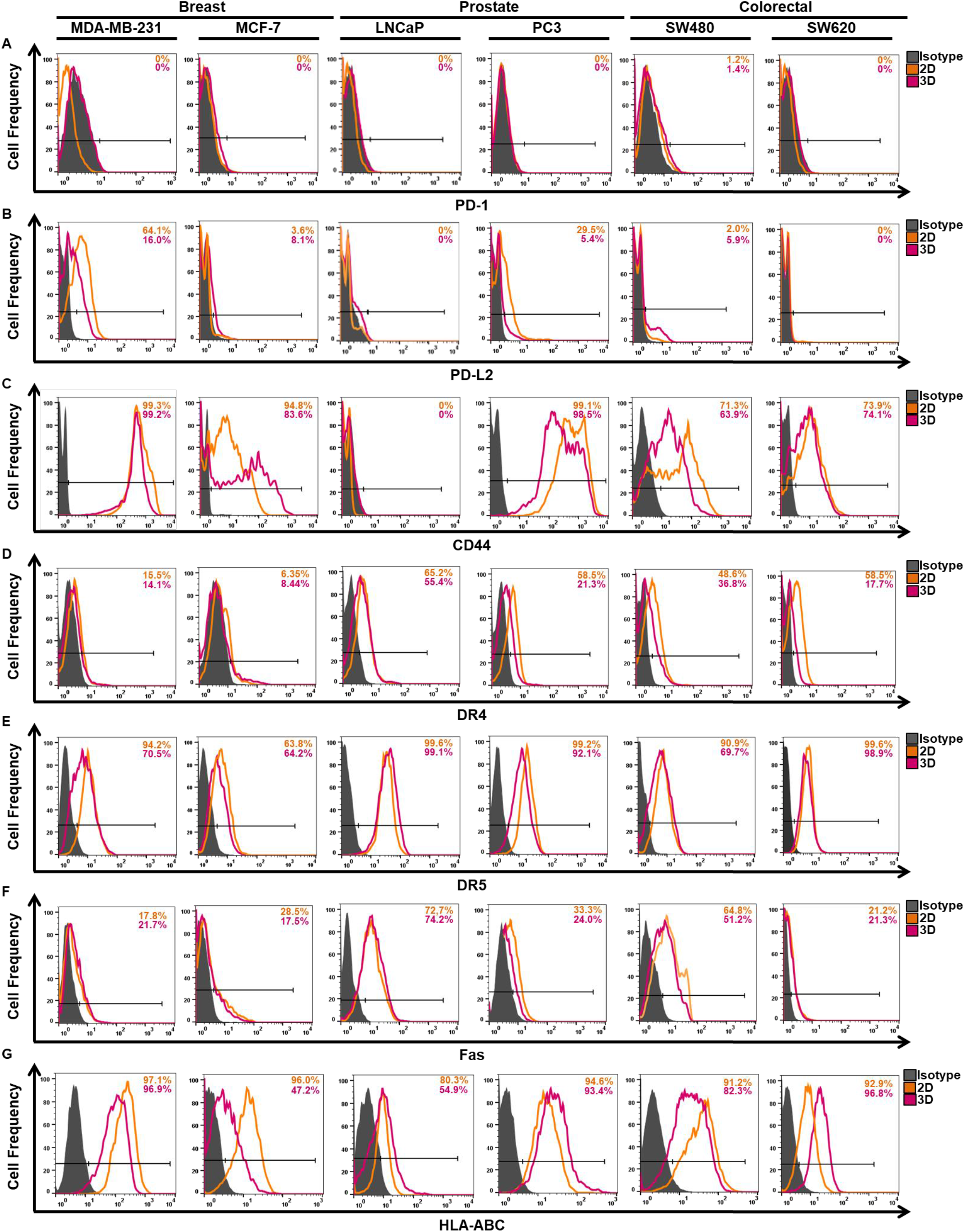
Representative flow cytometry histograms of different immunological markers expressed by human cancer cell lines grown in 3D spheroids formed by hanging drop compared to cells grown in 2D cell culture. Representative flow cytometry histograms are displayed for breast, prostate and colorectal cancer cell lines of different immunological markers, including PD-1 (**A**), PD-L2 (**B**), CD44 (**C**), DR4 (**D**), DR5 (**E**), Fas (**F**) and HLA-ABC (**G**) measured in 2D and 3D culture to show changes, if any, in the level of protein expressed and the average percentage of cells expressing each protein (normalised to the isotype control). 2D/3D isotype control (grey), 2D-cultured cells (orange) and hanging drop 3D spheroids (pink). Each histogram is representative of n=3 independent experiments.

**Figure 9.**
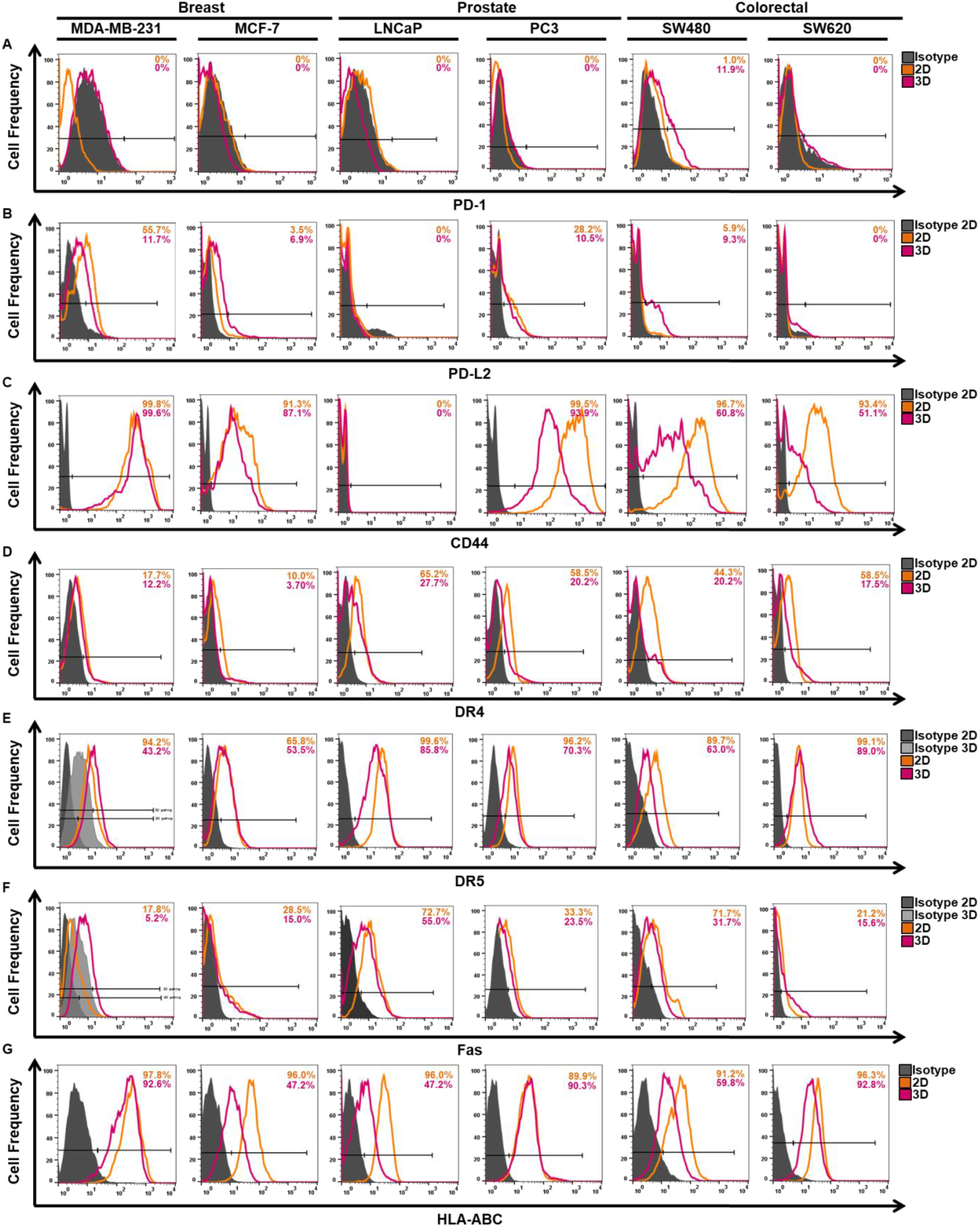
Representative flow cytometry histograms of different immunological markers expressed by human cancer cell lines grown in 3D spheroid colonies generated in alginate compared to 2D-cultured cells. Representative flow cytometry histograms are displayed for breast, prostate and colorectal cancer cell lines of different immunological markers, including PD-1 (**A**), PD-L2 (**B**), CD44 (**C**), DR4 (**D**), DR5 (**E**), Fas (**F**) and HLA-ABC (**G**) measured in 2D and 3D culture to show changes, if any, in the level of protein expressed and the average percentage of cells expressing each protein (normalised to the isotype control). 2D/3D isotype control (grey), 2D-cultured cells (orange) and alginate 3D spheroid colonies (pink). Each histogram is representative of n=3 independent experiments.

**Figure 10.**
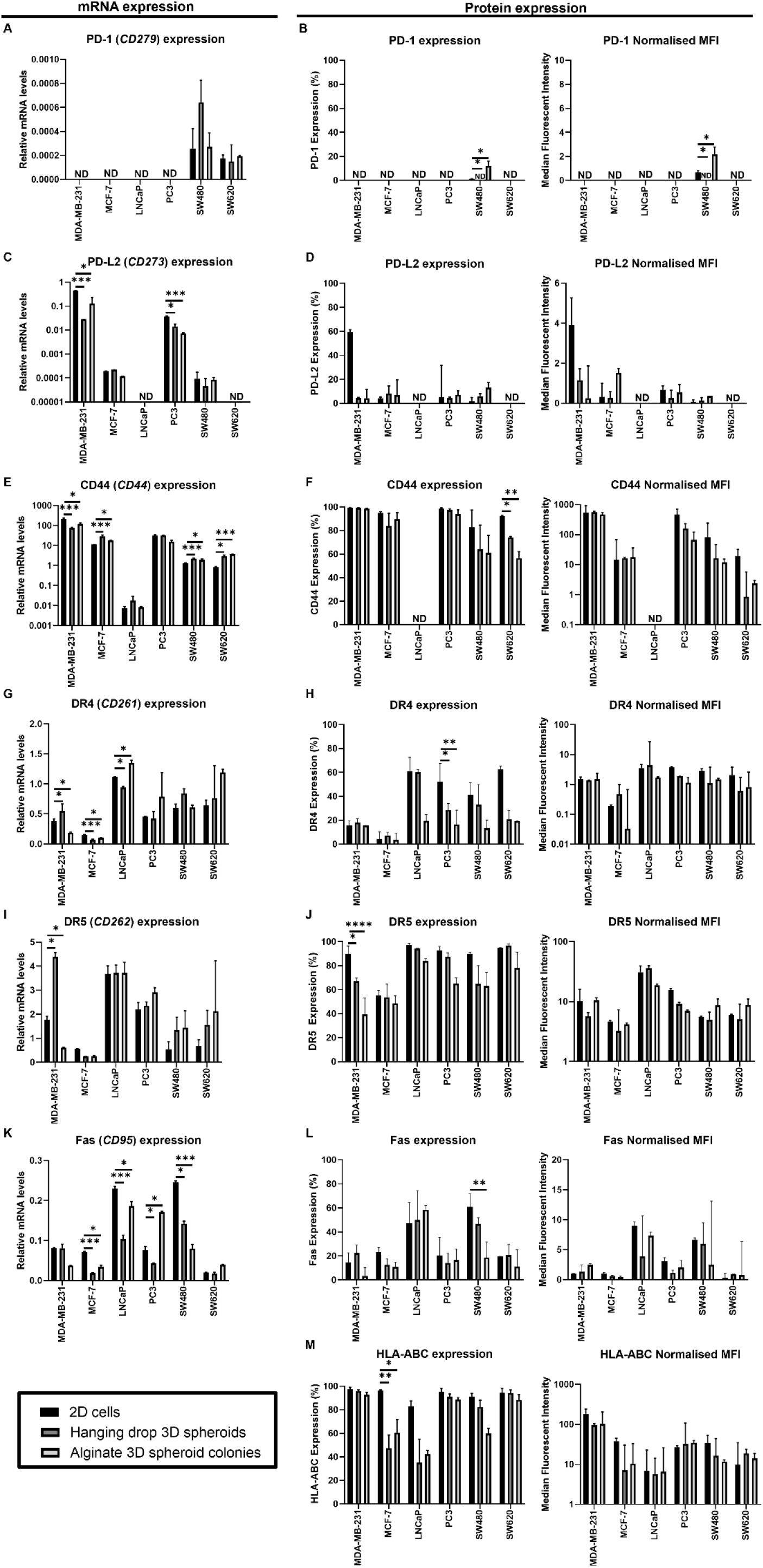
The expression of immunological markers (PD-1, PD-L2, CD44, DR4, DR5, Fas and HLA-ABC) at mRNA and/or protein level by breast, prostate and colorectal cancer cells grown in a 3D cell culture models compared to their 2D counterparts. qRT-PCR and flow cytometry were used to measure the mRNA and protein levels of different immunological markers including: PD-1 (*CD279*) (**A, B**), PD-L2 (*CD273*) (**C, D**), CD44 (*CD44*) (**E, F**), DR4 (*CD261*) (**G, H**), DR5 (*CD262*) (**I, J**), Fas (*CD95*) (**K, L**) and HLA-ABC (**M**) expressed by human breast, prostate and colorectal cancer cell lines grown in 3D cell culture models compared to cells grown in 2D monolayer culture. Expression data was acquired at day 3 for hanging drop 3D spheroids and at day 10 for alginate 3D spheroid colonies. Bars on each graph represent: 2D-cultured cells (black), hanging drop 3D spheroids (dark grey) and alginate 3D spheroid colonies (light grey) for each cell line. Protein expression data is shown has the percentage of cells expressing each immunological marker, together with the normalised MFI for each immunological marker. Data is presented as median ± IQR, n=3 independent experiments each with 3 technical repeats, Kruskal-Wallis followed by Conover-Iman multiple comparisons test, *P<0.05, **P<0.01, ***P<0.001, ****P<0.0001, ND, non-detectable.

To further characterise cancer cells and determine their immunogenic status, we measured the expression of death receptors (DR4, DR5 and Fas) at mRNA and protein level and the level of major histocompatibility complex (MHC)-class 1 antigens (HLA-ABC) on the cell surface of cancer cells in a 3D environment compared to 2D. In 3D culture, MDA-MB-231, MCF-7 and LNCaP cells displayed significant changes to DR4 (Figure 10G) and/or DR5 (Figure 10I) mRNA expression compared to 2D. MDA-MB-231 and LNCaP cells demonstrated a different expression pattern depending on the 3D model in which the cancer cells were implemented in at mRNA level. However, at protein level these significant changes observed in 3D were reversed showing the opposite effect or similar levels to 2D-cultured cells. DR4 (Figure 10H) and DR5 (Figure 10J) protein expression in 3D cell culture by all cancer cells investigated was either similar or reduced compared to 2D cell culture. In PC3 and MDA-MB-231 cells, the reduced expression of DR4 and DR5 was significantly altered in both 3D models compared to 2D, respectively. Similarly, Fas mRNA (Figure 10K) and protein (Figure 10L) expression in 3D-cultured cells was either comparable or reduced in comparison to 2D-cultured cells in most cell lines investigated. MCF-7, LNCaP, PC3 and SW480 cells displayed a significant decrease in Fas expression at mRNA level in both 3D culture models. SW480 cells also displayed a significantly reduced frequency of cells expressing Fas in 3D culture models compared to 2D. Additionally, the proportion of cancer cells expressing HLA-ABC on their cell surface was reduced in 3D cultures compared to 2D-cultured cells for most cancer cells, though; this reduction in expression was only significant for MCF-7 cancer cells. The level of HLA-ABC expression was comparable across all cultures for all cell lines investigated.

This data, along with the PD-L1 expression data, demonstrates that by culturing cancer cells in a 3D cell culture environment compared to standard 2D cell culture influences different cancer cell phenotypes, here, specifically associated with immunological marker expression that may be more comparable to their expression in an *in vivo* human tumour to allow for improved pre-clinical *in vitro* investigations.

## Discussion

There is currently heightened interest in understanding the role of PD-L1 in the tumour and the tumour microenvironment, since targeting the PD-1/PD-L1 signalling axis has revolutionised the cancer therapy landscape, yet challenges remain, and need addressing to improve patient response and overcome resistance mechanisms in all cancers. Over recent years 3D cell culture models have been reported to have more physiological relevant functions for oncology and immuno-oncology studies than standard 2D cell culture in terms of defence response to treatment and immune system modulation. Additionally, whilst the use of *in vivo* mouse models is paramount to testing the efficacy and systemic effects of therapeutic drugs before human clinical trials, unless humanised mouse models are utilised, they are problematic for investigating the PD-1/PD-L1 signalling axis in a human context and hence for the testing of human anti-PD-1/PD-L1 monoclonal antibody therapies. Indeed, the expression of immune checkpoint molecules by tumours *in vivo* have been reported to differ from their expression in 2D cell culture models (Rom-Jurek *et al*., 2018; Boucherit *et al*., 2020), highlighting the importance of utilising more robust human 3D cell culture models to mimic characteristics of the tumour more closely to measure immunotherapy and tumour responses (Hudson *et al*., 2020). To this end, here we investigated whether the basal expression of PD-L1 by six human cancer cell lines could be modulated at protein level and whether its expression would be altered when grown in two different 3D cell culture models of varying *in vitro* complexity as opposed to standard 2D cell culture. We found that basal levels of PD-L1 expression by cancer cells could be modulated by cytokine treatment and that PD-L1 expression was altered in a 3D environment compared to 2D-cultured cells. Additionally, the expression of other immunological markers (PD-1, PD-L2, CD44, DR4, DR5, Fas and HLA-ABC) were found to change in cancer cells being grown in 3D cell culture models as opposed to 2D monolayer culture.

The basal expression levels of PD-L1 by human cancer cell lines were found to be in accordance with the published literature, in that, MDA-MB-231 breast and PC3 prostate cancer cells expressed high levels of PD-L1; LNCaP prostate and SW480 colorectal cancer cells expressed low to moderate levels of PD-L1; and MCF-7 breast and SW620 colorectal cancer cells expressed low levels of PD-L1 (Song *et al*., 2013; Soliman *et al*., 2014; Martin *et al*., 2015). Cytokine driven PD-L1 upregulation on the surface of cancer cells is a well-documented mechanism that occurs in the tumour microenvironment during the development of the adaptive immune response to prevent cancer cells being recognised and destroyed by tumour-specific CD8+ T cells (Pardoll, 2012). Our data confirms that human breast, prostate and colorectal cancer cells investigated here have the potential to demonstrate adaptive immune resistance. Pro-inflammatory cytokines, IFNγ and TNFα, were able to individually increase cell surface PD-L1 expression on cancer cells and work synergistically in some cancer cells to further increase cell surface PD-L1 expression; a synergistic mechanism only previously observed in human dermal lymphatic endothelial cells (Yee *et al*., 2017). Whilst 2D cell culture provides opportunities to explore the biological functions of cancer cells in an easy and high throughput manner, cancer cells display aberrant characteristics when grown in monolayer, which fails to mimic those observed in *in vivo* human tumours. Many studies have shown that PD-L1 expression by human cancer cells differs *in vivo* compared to their *in vitro* monolayer counterparts (Gatalica *et al*., 2014; Rom-Jurek *et al*., 2018). In the present study, we utilised a scaffold-free (hanging drop method) and a scaffold-based (alginate hydrogel bead) system to form 3D spheroids and 3D spheroid colonies, respectively, to assess PD-L1 expression by human cancer cells compared to their 2D monolayer counterparts. These 3D cell culture models have previously been reported to sustain an oxygen and nutrient gradient (Muller-Klieser and Sutherland, 1982; Alessandri *et al*., 2013; Nunes *et al*., 2019), encourage increased extracellular matrix deposition (Lee and Mooney, 2012; Rios de la Rosa *et al*., 2018), facilitate genomic and protein alterations (Souza *et al*., 2017; Souza *et al*., 2018) and demonstrate increased resistance to anti-cancer therapies (Luca *et al*., 2013; Riedl *et al*., 2017; Takahashi *et al*., 2020; Boucherit *et al*., 2020), similar to that observed in *in vivo* human tumours. Here, we show new data that PD-L1 mRNA expression was reduced in breast, prostate and colorectal cancer cells grown in both 3D cell culture models compared to their 2D monolayer counterparts. This reduced PD-L1 expression in 3D culture was also observed on the cell surface of MDA-MB-231 breast and PC3 prostate cancer cells. It is well documented that TNBCs have the highest PD-L1 expression among all breast cancer subtypes (Solliman *et al*., 2014; Gatalica *et al*., 2014). Consistent with this, MDA-MB-231 TNBC cells in 3D cultures still expressed the highest level of PD-L1 compared to MCF-7 luminal A breast cancer 3D cultures, despite their reduced expression. A study analysing PD-L1 expression on human tumour samples demonstrated that 59% of TNBCs expressed PD-L1 compared to 33% of the luminal A subtypes (Gatalica *et al*., 2014). In the same study, out of 20 prostate and 87 colon tumour samples only 25% and 21% were shown to express PD-L1, respectively. This study indicates that PD-L1 is heterogeneously expressed among human cancers *in vivo* and that PD-L1 expression is only found in a small percentage of tumour samples. Martin *et al*., (2015) also demonstrated the paucity of PD-L1 expression in human prostate cancer cell lines (including LNCaP and PC3 cancer cells) and human prostate samples to find that PD-L1 was expressed at very low levels across all samples (Martin *et al*., 2015). In line with our study, it was recently reported that human breast cancer cell lines (BT-474, MDA-MB-231, SK-BR-3, and JIMT-1) that had been implemented into immunodeficient and humanized mouse models developed tumours that displayed diminished PD-L1 expression compared to their 2D *in vitro* counterparts (Rom-Jerek *et al*., 2018). They also found that PD-L1 was not expressed ubiquitously across the tumour. Similarly in *in vivo* human tumours not all cancer cells show uniform PD-L1 expression and their PD-L1 status is likely to vary from any one time given the influence of the tumour microenvironment (Tumeh *et al*., 2014; Ribas and Hu-Lieskovan, 2016). The downregulation of PD-L1 expression by cancer cells *in vivo* was thought to be associated with the cell density and compactness of the tumour, and the lower proliferative rate of the cancer cells, as opposed to the aberrantly reduced cell-cell contact and higher proliferative rate cancer cells exhibit in 2D culture, respectively (Satelli *et al*., 2016; Clark *et al*., 2016; Rom-Jerek *et al*., 2018). With the increased cell-cell contact and the reduced proliferative status of cancer cells that is observed in 3D cell culture, we can postulate that this may in part explain our observations and also implies that our 3D cancer models may be able to recapitulate PD-L1 expression to that of an *in vivo* tumour for the first time. To further support our data, more recently, substrate stiffness has been reported to modulate PD-L1 expression in lung (Miyazawa *et al*., 2018) and breast (Azadi *et al*., 2019) cancer cells, highlighting how mechanical cues in the tumour microenvironment can also influence PD-L1 expression. These studies showed that softer substrates facilitated a decrease in PD-L1 expression. Therefore, considering monolayer cells are cultured on hard plastic with stiffness in the gigapascal range which is not comparable to human soft tissues, either healthy or pathological, therefore it is not unexpected that in 3D cultures where cancer cells are in suspension (hanging drop 3D spheroids, elastic moduli <0.1Kilopascal, KPa) or in 1.2% alginate (3D spheroid colonies, elastic moduli of ∼5-10KPa) that PD-L1 expression was reduced, where the stiffness is more physiologically relevant to *in vivo* tumour tissue.

Although cancer cells in this study present with reduced PD-L1 expression at mRNA level in a 3D culture environment compared to their 2D counterparts, some cancer cells including MCF-7 breast, LNCaP prostate, SW480 and SW620 colorectal cancer cells displayed an increased proportion of cells expressing cell surface PD-L1 at a higher MFI in 3D in at least one or both of the 3D cell culture models compared to 2D-cultured cells. This could suggest that other intrinsic and extrinsic factors that regulate PD-L1 are playing a role in the 3D environment to promote PD-L1 protein expression on a higher proportion of the cancer cells (Hudson *et al*., 2020). Within the literature, only one previous publication has investigated PD-L1 expression in 2D versus 3D cell culture (Lanuza *et al*., 2018). In this study, only colorectal cancer cell lines (HCT116, HT29 and Caco-2) were assessed; none of which we studied here and only one 3D model was used. They showed HT26 and Caco-2 colorectal cancer cells displayed increased MFI of PD-L1 expression in 3D cell culture, whilst the HCT116 displayed similar MFI to 2D-cultured cells. These results, along with ours, may suggest that PD-L1 regulation in a 3D environment is dependent on the cancer cells being implemented and highlights the heterogeneity among different cancer types. Many factors that have been shown to positively correlate with PD-L1 expression in native human tumours exist in 3D spheroids and potentially could account for the increases in cell surface PD-L1 we observe in this study. Some of these include: increased expression of HIFs (HIF-1α and HIF-2α) (Noman *et al*., 2014; Barsoum *et al*., 2014; Scharping *et al*., 2017); increased expression of GLUT-1 (Young *et al*., 2011); increased activation of oncogenic signalling pathways such as phosphatidylinositol 3-kinase (PI3K)/AKT and mitogen-activated protein kinase (MAPK) pathways (Riedl *et al*., 2017; Dong *et al*., 2018) and increased pro-inflammatory cytokine secretion such as IFN-γ and interleukin 10 (Mahon *et al*., 2015). Alternatively, the prevalence of PD-L1 expression in patient tumours has been reported to be higher in metastatic tumours compared primary tumours in some cancers (Wang *et al*., 2017). In 3D culture, MCF-7, LNCaP and SW620 cancer cells derived from metastatic tumours displayed increased PD-L1 protein expression which could potentially reflect their metastatic phenotype *in vivo*. However, SW480 cancer cells derived from a primary tumour also displayed increased PD-L1 protein expression in 3D culture to similar levels of its metastatic counterpart SW620 cancer cells. This once again highlights tumour heterogeneity and demonstrates how PD-L1 expression can vary, particularly amongst primary and metastatic tumours. In some cancers including breast (Tawfik *et al*., 2018) and urothelial carcinomas (Burgess *et al*., 2019), primary- and secondary-derived tumours have been shown to express similar levels of PD-L1. In colorectal cancer specifically, the status of PD-L1 positivity and how this reflects the tumour stage remains to be elucidated.

Whilst a consistency in the expression pattern of PD-L1 was observed across both 3D cell culture models, this was not the case for the other immunological markers investigated. Indeed, the different 3D cancer models used have distinctive characteristics in that the hanging drop method facilitates heterogeneous aggregation of cancer cells (Knight and Przybors, 2015), whereas the alginate hydrogel beads simulate the formation of clonal spheroids, whereby cancer cells are being selected for survival characteristics and their ability to self-renew and proliferate from a single cell (Florczyk *et al*., 2016). Recently, transcription factor NRF2 has been shown to be a prerequisite for clonal formation in order to protect cancer cells from oxidative stress that develops during tumorigenesis (Takahashi *et al*., 2020). If NRF2 expressing cells are being selected for in the alginate model, this could subsequently alter the level of genes and proteins expressed, and therefore may be responsible for the differences we observe in this study when comparing the expression data from the alginate model to the 3D hanging drop model and 2D cell culture. The changes in the expression of PD-1 protein by colorectal SW480 cells and in the expression of DR4 and DR5 mRNA expression by MDA-MB-231 breast and LNCaP prostate cells in this study illustrate how different 3D models can influence different cancer cell characteristics depending on the cancer type. In the alginate model, PD-1 was upregulated on the surface of SW480 cells compared to 2D-cultured cells and 3D spheroids. Conversely, DR4 and DR5 were downregulated by MDA-MB-231 cells in the alginate model whereas they were upregulated in the hanging drop model compared to 2D-cultured cells. Collectively, these findings could indicate that these specific cancer cell lines are being selected for clonal expansion in the alginate model to ultimately increase the expression of genes and proteins that would likely be advantageous for cancer cell survival (i.e. tumour-PD-1 expression could intrinsically promote cancer cell survival (Hudson *et al*., 2020)), as well as decrease the expression of immunological markers that would otherwise increase their susceptibility to immune-mediated cell death in the tumour microenvironment (i.e. downregulating death receptors could make cancer cells less susceptible to immune-mediated killing and resistant to drugs that target death receptors (Shin *et al*., 2001; Matínez-Lostao *et al*., 2015; Chandrasekaran *et al*., 2014)). Reduced expression of death receptor Fas and HLA-ABC was also observed in our study, suggesting the possibility that cancer cells have the tendency to reduce their immunogenic status when cultured in a 3D environment which more closely mimics that of an *in vivo* human tumour (Ashkenazi *et al*., 2008; de Carvalho-Neto *et al*., 2013; Peter *et al*., 2015; Dhatchinamoorthy *et al*., 2021). In support of this statement, Chandrasekaran *et al*., (2014) demonstrated how breast cancer (BT20 and MCF-7) 3D tumour spheroids were more resistant to TNF-alpha-related-apoptosis-inducing-ligand (TRAIL)-mediated apoptosis than 2D-cultured cells due to their downregulation of DR4 and DR5. Restoring DR4 and DR5 expression via COX-2 inhibition increased TRAIL-mediated apoptosis in 3D cultures (Chandrasekaran *et al*., 2014). In the same study, cells in 3D culture were shown to exhibit high CD44 expression, a cancer stem cell-like characteristic that facilitates tumorigenic processes such as proliferation, invasion and metastasis (Senbanjo and Chellaiah, 2017). In our study 4 out of the 6 cancer cell lines upregulate CD44 mRNA in 3D cell culture models compared to 2D which once again highlights the need for using 3D cell culture over 2D cell culture to mimic *in vivo* tumour characteristics.

In summary, there are several key regulators of PD-L1 expression that have previously been reported (Hudson *et al*., 2020) and many of these regulators can be influenced by growing cancer cells in 3D (Riedl *et al*., 2017; Souza *et al*., 2018), and thus may contribute to the changes in PD-L1 observed in this study. Further investigations are required to determine the exact mechanism of PD-L1 up- or down-regulation in each cancer cell line in 3D, compared to conventional 2D monolayer. Importantly, the alterations to PD-L1 expression by cancer cells that we observed were consistent across two different 3D cell culture models, which suggests that the mechanisms for regulating PD-L1 expression in 3D cell culture is intrinsic to these cancer cells. Both 3D models explored in this study present with advantages over 2D monolayer culture, allowing better mimicking of *in vivo* conditions for the investigation of PD-L1. The further characterisation performed in this study to assess other immunological markers expressed by human cancer cells in 3D cell culture models compared to 2D-cultured cells provides a platform for future oncology and immuno-oncology studies investigating anti-cancer therapies which may target these markers either alone or in combination with PD-L1-targeted therapy.

## Conclusion

PD-L1 expression in solid tumours mediates immune evasion, drug resistance and tumour progression and is associated with poor prognosis. Immunotherapies targeting the PD-1/PD-L1 signalling axis are insufficient to reject all tumour types and therefore only benefit patient minority. To improve the clinical efficacy and better understand the mechanism of action of PD-L1 in tumour and the tumour microenvironment, culturing of human cancer cells in 3D is necessary to recapitulate *in vivo* tumour characteristics that are non-existent in 2D monolayer culture. Here, we have shown for the first time in these human cancer cell lines that PD-L1 expression is altered in 3D cell culture systems compared to 2D monolayer culture. Additionally, we have further characterised 3D cell culture models for their expression of other important immunological markers. These results emphasise the importance of using 3D cell culture has the level of expression of PD-L1 and other immunological markers by cancer cells in 3D is more likely to mimic that of an *in vivo* human tumour than standard 2D cell culture. Utilising 3D cell culture in this context may better able the investigation of the tumour-intrinsic role of PD-L1, treatment response to PD-1/PD-L1-targeted therapies and cancer cell-immune cell interactions. This could ultimately provide a better understanding of the tumour intrinsic role of PD-L1 and its role in the tumour microenvironment and potentially could enhance the transferability of PD-1/PD-L1-targeted therapy and combination regimes into the clinic by bridging the gap between *in vitro* 2D monolayer and *in vivo* studies.

## Acknowledgements

The authors wish to thank the Biomolecular Sciences Research Centre and Sheffield Hallam University for funding this work.

KH conducted the experiments, analysed the data, wrote and revised the manuscript with support from RL, NC and NJM. KH, RL, NC and NJM designed the experiments. RL, NA and NJM designed the project. All authors provided feedback for the manuscript and approved the final version.

## Statements and Declarations

The authors have no relevant financial or non-financial interests to disclose.

